# Nanofluidic Device for Manipulation and Modification of DNA by Proteins

**DOI:** 10.1101/2022.12.29.521498

**Authors:** Saroj Dangi, Ming Liu, Zubair Azad, Preston Countryman, Maedeh Roushan, Gideon I. Livshits, Parminder Kaur, Hai Pan, Zhubing Shi, Ariana C. Detwiler, Patricia L. Opresko, Hongtao Yu, Hong Wang, Robert Riehn

## Abstract

Single-molecule techniques provide important details supplementing the framework obtained from traditional bulk experiments. Many cellular processes such as DNA replication, DNA repair, and telomere maintenance involve interaction among multiple proteins, their co-factors, and DNA. To investigate such interactions and to differentiate the function of each component necessitate a technique that allows the combinatorial exposure of DNA to multiple proteins and co-factors as well as manipulation of the DNA configuration. We present a nanofluidic device with the capability of active combinatorial exchange of up to three buffers in real-time and dynamic manipulation of DNA under physiologically relevant conditions. We demonstrate its utility in monitoring compaction of DNA by telomeric proteins, DNA modification by an endonuclease, and DNA loop extrusion by cohesin.

## I. INTRODUCTION

A multitude of single-molecule techniques that allow the visualization and manipulation of individual DNA have been explored^1–7^. The capability of single molecule techniques to observe the trajectories of DNA conformations is crucial in the study of DNA-protein interactions, which often cannot be captured using bulk experiments because of the large variability of possible conformations. Despite the rapid development enabling the manipulation of individual DNA, active exchange of protein-carrying buffers *in-situ* has in most cases been based on the tethering of DNA ends^8,9^. Thus, we have to presume that the activity of proteins and the dynamics of protein-DNA interactions carry not only information intrinsic to the protein-DNA interaction, but also the tension along the DNA molecule^10^. Here we present a tension-free technique which allows manipulation of genomic-sized DNA under physiologically relevant conditions. It allows fluorescence microscopy observation, the dynamic co-location of multiple DNA molecules, exposure of DNA to two proteins with arbitrary exposure sequence, observation of covalent DNA modification, and DNA loop extrusion by a protein.

Our focus here is on techniques that are based on the concept of stretching DNA by confining it to a quasi-1d channel^7^, which is typically referred to as nanochannel stretching or nanofluidic stretching of DNA. The physics of stretching DNA have been explored at considerable depth^11–19^. Stretching in narrow channels, say channels with a square cross-section of less than 50×50 nm^2^, follows the stiffness-dominated Odijk regime that is rather insensitive to DNA-DNA self-interactions, and is commonly employed in sequence-mapping applications. In contrast, stretching in wider channels, say with a cross-section of 150×150 nm^2^, follows the extend de Gennes regime that is fundamentally based on self-avoidance of polymer chains. Stretching in wide channels also renders the DNA fundamentally soft in the sense that the magnitude of short-range density fluctuations is comparable to the local density of DNA^20–25^, which approaches a regime that is similar to the nuclear environment. The combination of the freedom of DNA to change its conformation due to thermal fluctuations, the sensitivity of the stretching to DNA-DNA interactions, and the high local density make nanochannel-stretched DNA an ideal test-bed for protein-induced conformational transitions that can yield statistically significant results even if only a modest number of molecules are observed. Importantly, the stretching is independent of substrate length, the conformations can be monitored continuously using fluorescence microscopy, and there is no axial tension along the DNA molecule. The absence of tension is in particular important since flow-based assays can introduce a position dependence^26^.

Devices based on quasi one-dimensional layouts have been successful in mapping the binding location of proteins as markers of transcriptional or epigenetic states^27–32^ and monitoring the changes to DNA conformation induced by protein binding^33–47^. Earliest attempts to capture time-dependent activity of proteins used reaction-diffusion gradients of protein co-factors such as magnesium^48^, or diffusion of the proteins themselves^37^. However, that limits the active scale size of a device or the time resolution.

The limitation imposed by the diffusion speed can be removed if the buffer containing protein or protein co-factors can actively be moved through the device at a different velocity than the DNA. An early pathway to decouple polymer motion and buffer flow was based on electrokinetic effects and non-uniform channel geometries^11,49^. While the electrokinetic effects lead to a real-time tunability of potentials, they are notoriously complex under nanoconfinement^50^ and impose a narrow range of feasible buffer conditions. An alternate approach relies on junctions of nanochannels, either individually controlled^51^ or as part of an engineered lattice^52^, which in principle is a powerful approach but is sensitive to single-channel blockages that render the entire device non-functional.

The most promising concept, however, is to employ a confinement energy landscape that is established by a nanoslit whose surface is nano-patterned with nanogrooves, or other lateral nanoscale features^53**?**, 54^. We call this concept mixed 2d-1d-confinement here. Through tuning of the height of the slit-like confinement and the size of the lateral confinement of the grooves, the equilibrium configuration can be shifted towards long DNA residing entirely within the grooves. DNA that is confined to the grooves behaves as if it were confined to a nanochannel. The key feature for the study of protein-DNA interactions is that the buffer and the DNA now have different transport properties^55–57^ that allow for a buffer exchange. The concept was used in a considerable number of investigations of protein-DNA interactions that report conformational and topological modifications of DNA^58–60^. The most promising concept, however, is to employ a confinement energy landscape that is established by a nanoslit whose surface is nano-patterned with nanogrooves, or other lateral nanoscale features^53**?**, 54^. We call this concept mixed 2d-1d-confinement here. Through tuning of the height of the slit-like confinement and the size of the lateral confinement of the grooves, the equilibrium configuration can be shifted towards long DNA residing entirely within the grooves. DNA that is confined to the grooves behaves as if it were confined to a nanochannel. The key feature for the study of protein-DNA interactions is that the buffer and the DNA now have different transport properties^55–57^ that allow for a buffer exchange. The concept was used in a considerable number of investigations of protein-DNA interactions that report conformational and topological modifications of DNA^58–60^.

Here we expand the range of application of the mixed 2d-1d confinement device paradigm. We show that the sequence of proteins to which a DNA molecule is exposed, which is controllable within the device, can have physiological impact on a DNA complex with multiple shelterin proteins. We further show that nanogroove geometries with junctions, such as Y-shaped channels, hold promise in the manipulation of DNA configurations, and that they allow the direct imaging of DNA loop extrusion by cohesin.

## II. MATERIALS AND METHODS

### A. Device Design

The layout of the nanofluidic device is illustrated in Fig. 1. The device has four pairs of feeding microchannels converging towards the center of the chip, where a reaction site with an area 100×100 µm^2^ is located. Each arm of the feeding microfluidic channels is about 12 mm long from the access port to the central reaction site, 80 µm wide, and 1 µm deep. The reaction site is connected to microchannels through nanochannels or nanoslits. The nanoslits are used for flowing in proteins or reagents to the reaction site, while the nanochannels are used for injecting DNA molecules to the reaction site and flushing away buffers after the reaction is complete. The channel dimensions are chosen such that the resistances of fluid flow from microchannels to the reaction site through either nanochannels or nanoslits are approximately equal. This enables us to balance the hydrodynamic stresses at the reaction site along all axes for finer control of DNA manipulation. In addition, the balance of fluid flow resistances minimizes the premixing of buffers during the flushing of microchannels.

**FIG. 1.**
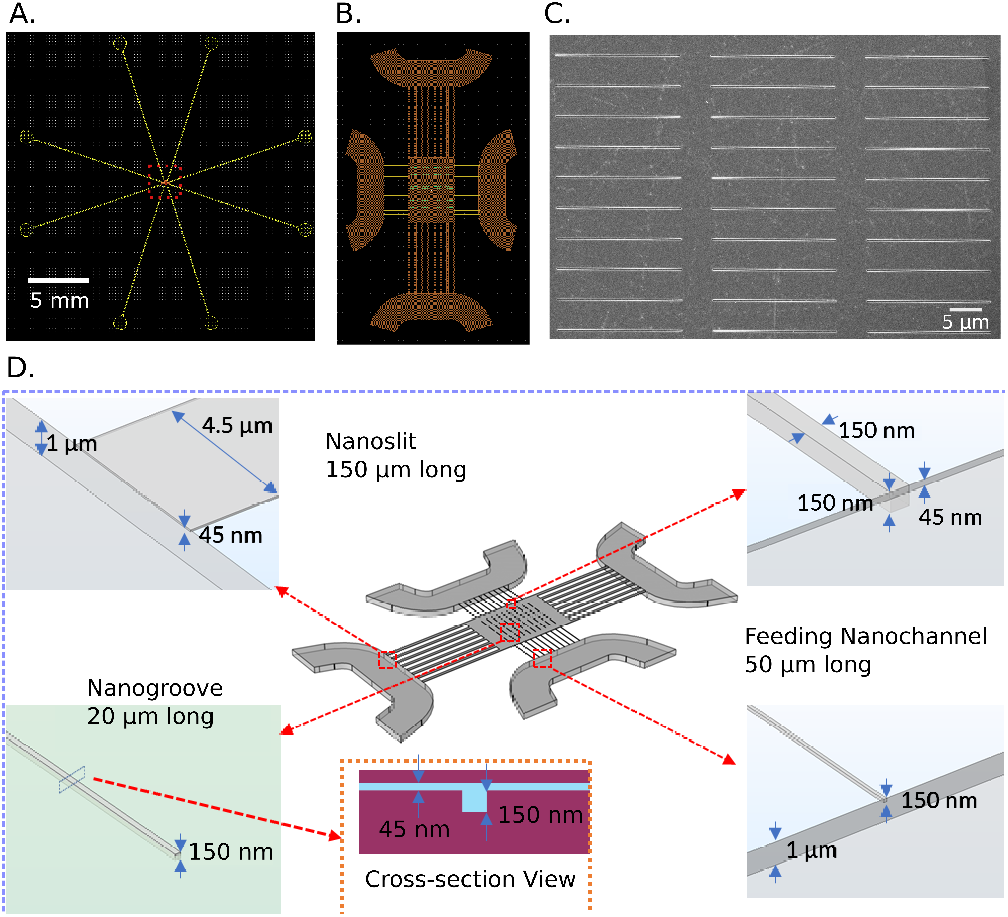
Nanofluidic device structure. **A**. Layout of the chip which has four pairs of access ports, each pair connected by a “U” shaped microchannel. The central reaction site is shown by a rectangle.**B**. Layout of the reaction region where the straight nanogrooves are arranged like a parking lot. **C**. Scanning electron micrograph of the nanofluidic region showing nanochannels. **D**. Schematic of the nanofluidic region showing the dimension of channels. There are 8 nanoslit channels on each side of the reaction site. The nanoslits are 150 µm long with a cross-section of 4.5 µm × 45 nm (width × depth). The nanochannels are 50 µm-long with a cross-section of 150×150 nm^2^. At the center, the reaction site with an area of 100×100 µm^2^ contains the nanogrooves. The cross-sectional view of the 150×150 nm^2^ nanogroove is shown in the inset.

The reaction site is formed by a 45 nm-high nanoslit into which nanogrooves with a cross-section of 150×150 nm^2^ are recessed. We utilized two geometric layouts of the grooves within the nanoslit. The first layout is an array of straight 20 µm-long nanogrooves that are arranged like a parking lot such that the groves are along the axis of the nanochannels that carry DNA and perpendicular to the axis of the nanoslits that carry protein solution. DNA molecules confined inside the groves become trapped, and active exchange of buffers is carried out by flowing protein-containing buffer over the confined DNA molecule. The second layout has arrays of Y-shaped channels with their stems either aligned along or perpendicular to the flow axis of the nanoslit.

### B. Device Fabrication

The devices were fabricated on 550 µm-thick fused silica substrates. First, the shallow nanoslit region was pattered using photolithography, and etched with reactive ion etching (RIE). Then, the nanochannels and nanogrooves were patterned using e-beam lithography, followed by RIE. Finally, microchannels and access ports were fabricated with photolithography and RIE. The channels were sealed with a 150 µm-thick coverslip using the thermal bonding.

### C. Flow Control and Data Analysis

At the beginning of the experiment, the four microchannels of the device (Fig. 2) are loaded with buffer carrying DNA, buffers containing generically-named “protein 1” and “protein 2”, and flushing buffer, respectively. The 8-port device is controlled through arbitrarily controllable pneumatic nitrogen gas pressures between ±35 psi with a precision of 0.1 psi at the location of the ports. A sequence of the experiment starts with an empty reaction region (Fig. 2A), into which DNA is injected (Fig. 2B). DNA molecules are randomly confined by the nanogrooves. Then, DNA is exposed to buffer carrying “protein 1” (Fig. 2C) followed by “protein 2” (Fig. 2D), or vice versa, through nanoslits that run perpendicular to the direction of the nanogrooves. Application of a negative pressure on one port of the inactive “protein” channel ensures that the DNA, the upstream inactive protein, and the flushing buffer in their respective microchannels do not become contaminated by the active “protein” buffer. In the next step, the reaction site is reset by flushing away DNA and proteins with a flushing buffer while protecting the upstream DNA and protein channels through application of negative pressure on the down-stream channels (Fig. 2E). Finally, fresh DNA and proteins are advanced within their microchannels through application of a positive pressure upstream (Fig. 2F). Characteristic pressures for introduction of DNA and flushing of channels is on the order of +15 psi. The characteristic pressure during exposure of DNA to protein is ±1 psi.

**FIG. 2.**
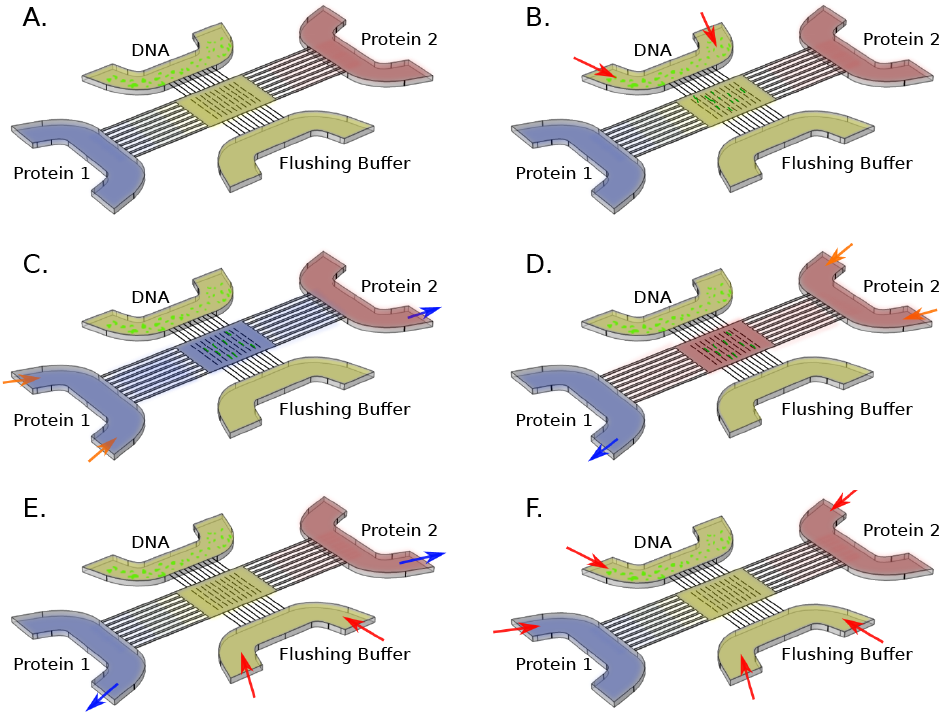
Procedure for performing sequential protein exposure of DNA with the nanofluidic device. Pneumatic driving of buffers with high positive (∼15 psi), low positive, and low negative (up to ± 1.0 psi) pressure are represented by red, orange, and blue arrows, respectively. **A**. Before actuation, and state during equilibrium observation. **B**. Injection of DNA. **C**. Injection of “protein 1”. **D**. Injection of “protein 2”. **E**. Removal of DNA from reaction zone. **F**. Flushing of feeding channels.

Throughout the process, the fluorescence signals from DNA and tracer dyes were monitored continuously using an inverted fluorescence microscope (Nikon) with a 60× Plan Apochromat objective (1.4 N.A.) or 100× Apo TIRF objective (N.A.=1.49), an IXon888 EMCCD (Andor), and a QV2 image splitter (Photometrics), from which we utilized a green (525/40) and a red (600/37) channel. Alternating illumination using 488 nm and 561 nm laser light (Coherent) and prevention of speckle pattern using a galvanic scanner (Cambridge Technology) allow independent quantification of emission from two chemical species.

### D. Biological Materials and Device Preparation

The majority of experiments were performed using 48.5 kbp-long linearized *λ* -DNA purchased from Promega that was stained using YOYO-1 (Invitrogen) at a molar ratio of 1 dye for every 10 basepairs. DNA was handled in 0.5× TBE buffer, unless otherwise noted.

For the demonstration of the impact of protein exposure sequences on protein activity, we chose TRF1 and TIN2, a pair of proteins involved in telomere maintenance^61^. A larger study involving an additional binding partner has been submitted for publication elsewhere^62^, and the biological significance is reported there. To mimic telomeric DNA, we ligated purified telomeric DNA after digestion of T270 plasmid (5.4 kbp, Addgene, PSXneo(T2AG3), #12403), which contains two telomeric sequence regions (TTAGGG)_135_ linked by a 23-bp nontelomeric sequence^63^. Only DNA molecules 20 kbp or longer were included in the data set. N-terminal His_6_-tagged TRF1 and N-terminal HA-tagged TIN2 (1-354 aa) were expressed in insect cells using the pFastBac system, as described previously^64–66^. Device passivation was performed using a physisorbed polyethylene glycol (PEG) on a dopamine adhesion layer^31^. The buffer used for the telomeric system was 20 mM HEPES pH 7.5, 100 mM KCl, 0.1 mM MgCl_2_.

For an experiment demonstrating restriction endonuclease activity, methyl cytosine-free *λ* -DNA was stained with YOYO-1 dye at 1:20 (dye: base pairs) ratio. The device was passivated using bovine serum albumine (BSA). DNA was loaded with a buffer containing 50 mM of potassium acetate, 20 mM of Tris-acetate, and 100 µg/ml of BSA at pH 7.7. SmaI (New England Biolabs) at the concentration of 1 U/ml was loaded along with DNA. The reaction was triggered by a buffer containing 4 mM of magnesium acetate in addition to the components listed above.

For the observation of ATP-dependent DNA loop extrusion, we used T4 phage DNA (169 kbp, FUJIFILM Wako) as our DNA substrate stained using YOYO-1 at a 1:20 dye to basepair ratio. The larger DNA length was necessary for ease of observation of looped and folded configurations. The cohesin complex, containing NIPBL as the DNA loader, as well as an ATPase-deficient mutant cohesin were prepared as previously described^67^. Cohesin-NIPBL and T4 phage DNA were incubated together for 5 min in a working buffer (20 mM Tris-HCl pH 7.5, 1 mM MgCl_2_, 50 mM NaCl) prior to introduction into the nanofluidic region to allow efficient loading. During loop extrusion, the buffer contained 2.5 mM ATP. The device was passivated in the same way as described for the telomeric protein system. Passive (flow-based) looping was demonstrated in the channel using the same DNA and buffers.

## III. RESULTS

This section is organized as follows: The functionality of the nanofluidic device for real-time active exchange of buffers with stationary DNA was tested using ionic strength gradients. That is followed by applications that utilize our specific design, namely the manipulation of DNA configurations by flow and shaped nanochannels, a demonstration that the protein exposure sequences in a binary protein system driving DNA compaction can dramatically impact the magnitude of compaction, and the demonstration the effect of two catalytically active protein systems.

### A. Verification of Device Layout

The real-time visualization of DNA extension in a buffer with changing ionic strength has been used as a device benchmark by other authors that study the time-resolved response of DNA to protein exposure^58,68^. The extension of DNA increases with a decrease in the ionic concentration of the buffer^69,70^. As a validation of our device layout and supporting instrumentation, we show the results of such an experiment in Fig. 3. YOYO-1 stained *λ* -DNA in 0.5× TBE buffer was injected to the reaction site, followed by exposure to 0.05× TBE buffer flowing from the “protein 1” side of the device (Figs. 3A and B). Switch-over points can be identified by dark lines, where we introduced a dark frame in the video by not activating the lasers. We observe an gentle increase of the end to end distance during the first 10 sec, followed by a plateau of the extension (Fig. 3C). After 25 sec, the flow was changed to expose the DNA to 5× TBE buffer from the “protein 2” channel to which Sulforhodamine 640 (here called rhodamine, 1 nM, observed under 561 nm illumination) had been added. A gradual decrease of the DNA extension was observed, together with an increase in the rhodamine fluorescence signal (Figs. 3C and D). At 50 sec, the flow was again changed to repeat the exposure to 0.05× TBE buffer, which resulted in an increase in DNA extension and a gradual decrease of the rhodamine signal (Figs. 3C and D).

**FIG. 3.**
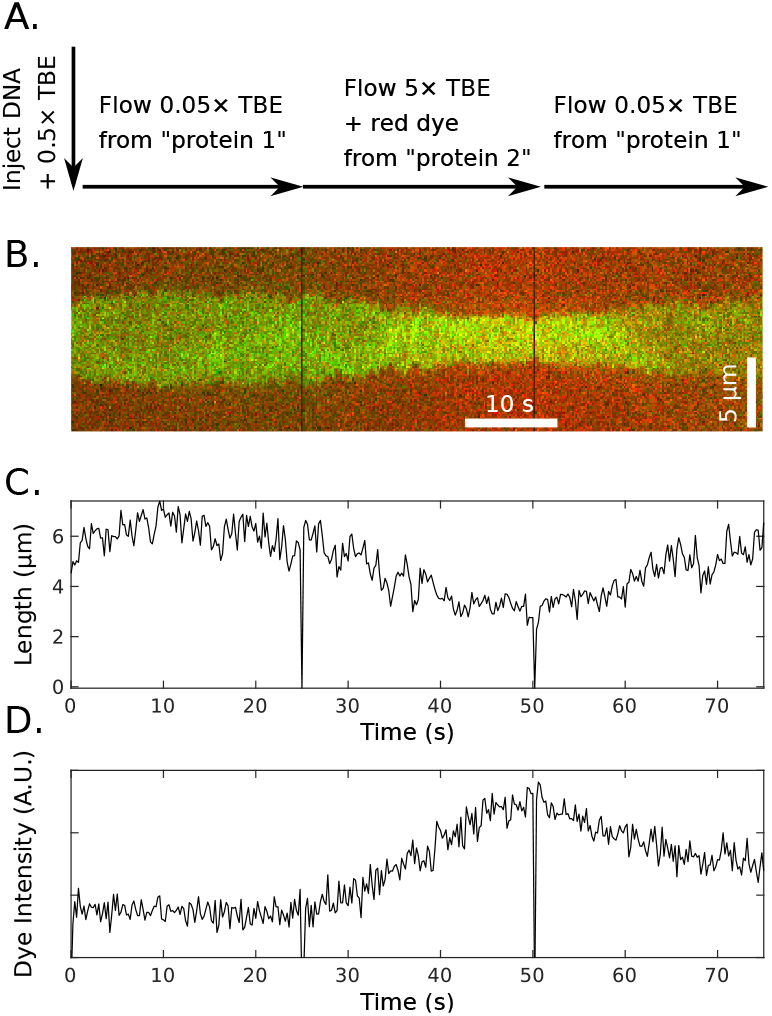
Validation of device using fluorescent dye and ionic strength-dependent DNA extension. **A**. DNA is injected in a 5× TBE buffer without rhodamine. It is then exposed to 0.05× TBE without rhodamine, followed by 5× TBE with rhodamine, and finally 0.05× TBE without rhodamine again. **B**. Kymograph showing the change in extension of DNA with varying buffer concentrations. The composite color image was obtained by merging green channel under 488 nm illumination (DNA) and the red channel under 561 nm illumination (rhodamine). The DNA center of mass was stabilized to allow for easy comparison. **C**. Graph of DNA extension versus time. **D**. Graph of average fluorescence intensity of red dye along channel versus time.

We quantified the image data of DNA extension and rhodamine fluorescence (Figs. 3C and D), leading us to the following observations. First, at the flow speeds employed by us in this experiment, the sharpness of the transition between two buffers and the transition speed are limited by the length of the nanoslit between the protein channels, the diffusivity of the ions and the dye, and the flow speed. A closed form of the time-dependence of the ion concentration under forced flow can in general not be given due to concentration polarization effects^71^, and so our discussion here is qualitative. Note in particular that the rhodamine signal started increasing almost immediately after the flow change at 25 sec, which indicates that a rhodamine gradient extends into the nanoslit at all times. The portion between 25 sec and 50 sec shows a sigmoid shape which suggests that the steady state rhodamine level was nearly obtained at 50 sec. However, we did not pursue longer exposures due to excessive oxidative DNA degradation in high salt. The diffusivity of the salt components should be higher than the diffusivity of rhodamine, and thus we anticipate that the same statements are true for the salt. Indeed, the extension contraction also shows a trend to an asymptote before 50 sec was reached. All protein systems in the following sections have lower diffusivities than rhodamine, and thus we anticipate sharper transitions for proteins.

### B. Dynamic Flow Manipulation of DNA

In a prior publication^51^, we had demonstrated the controlled folding or herniation of a single DNA molecule, as well as the co-location of two independent DNA molecules in a Y-shaped nanochannel. However, the capacity to perform these manipulations becomes only biologically relevant if it is possible to perform under exposure of proteins, and the closed quasi-infinite channels of the prior publication imposed strong constraints on the possible buffer flow rates while DNA was held in the field of view. In Fig. 4 we demonstrate that a modification of the pattern of nanogrooves within the nanoslit from straight to Y-shaped enables the present device to gain a similar functionality as our earlier device while allowing steady buffer flow over the molecule.

**FIG. 4.**
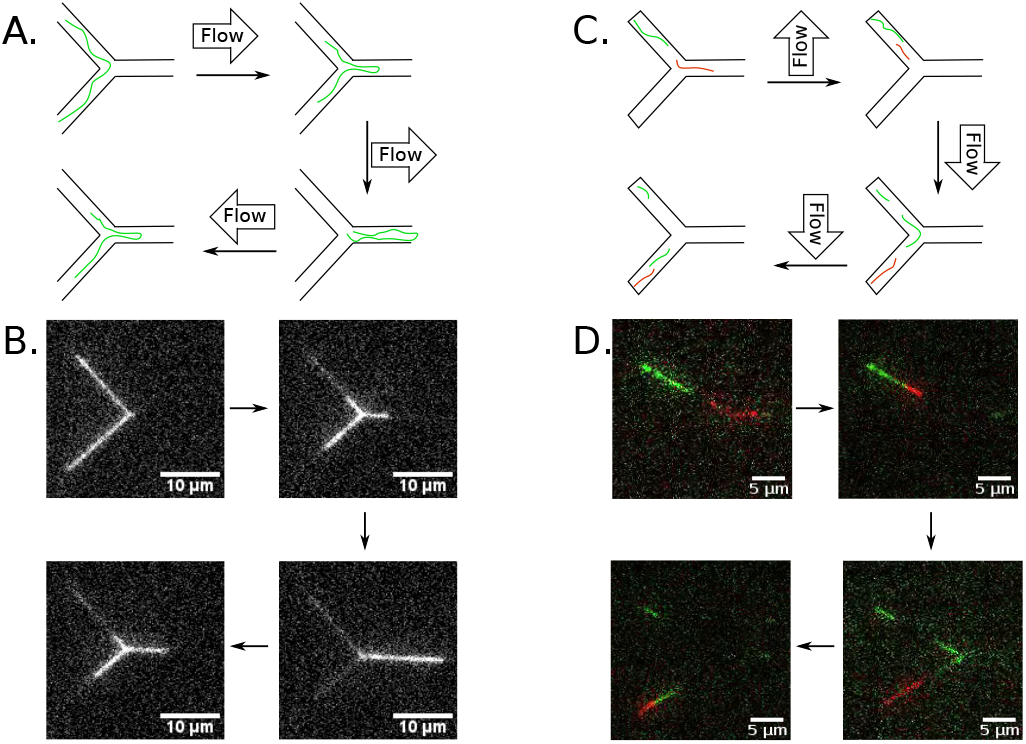
Flow-based manipulation of DNA in Y-shaped nanogrooves. **A**. Schematic of hernia extrusion by flowing along the symmetry axis of the Y. **B**. Snapshots of experimental realization of hernia extrusion. **C**. Schematic of co-locating the ends of two DNA molecules in flow perpendicular to the symmetry axis of the Y. **D**. Snapshots from experimental realization of co-location of ends of two DNA molecules.

The schematics in Fig. 4A and B demonstrate the principle of dynamic DNA configuration manipulations in flow. First we consider the folding or herniation of a single T4 DNA molecule at the 3-way junction formed by a Y-shaped channel (one stem and two branches) where the stem was parallel to the long axis of the nanoslit. DNA is initially brought into the nanoslit region and randomly located within a Y-shaped grove by a steady slow flow. The first manipulation step is to locate the molecule symmetrically in both Y-branches at the 3-way junction, either through a slow flow along the symmetry axis as described by Azad et al.^51^, or a flow perpendicular to it by flowing through the feeding nanochannels. The flow is then changed to be directed towards the stem of the Y, and a hernia is extruded within the stem. Once flow is stopped, the hernia spontaneously starts to unfold, but can actively be unrolled by reversing the flow direction.

For the second demonstration of flow-based DNA manipulation (Fig. 4C and D), YOYO-1 (green) and BOBO-3 (red) stained *λ* -DNA were pooled together to demonstrate overlap of two differently colored DNA molecules. The device differed from the one used for folding by having shorter Y-branches, and that the stem of the Y was perpendicular to the long axis of the nanoslit. As before, DNA is introduced into the nanoslit region of the device. Using slow flow, DNA are trapped in nanogrooves and randomly the same Y-shaped grove can be occupied by two differently-stained DNA. DNA can be forced into contact by flowing buffer perpendicular to the stem, which traps both DNA at the end of the branch channels with their end in close contact. In its ability to co-locate only the ends of DNA but not force a full overlap the device falls between the weak confinement reported by Capaldi et al. who showed co-location of two coil-like DNA molecules in one volume of a pit-and-nanoslit geometry^72^, and the extended overlap reported in a fully-enclosed Y-junction^51^. We believe that the ideas of leveraging differences in fluid and polymer transport under nanoconfinement49,52 could lead to striking DNA manipulation opportunities, such as location-dependent polymer velocities in undulating channels52.

### C. Compaction of DNA by a Multi-protein Complex

The monitoring of DNA configurations under impact of non-catalytic protein binding has developed into a key capability of nanofluidic analysis^33–47^. In particular, nanofluidic analysis is a sensitive probe of changes in the polymer physics parameters of effective width, persistence length as well as local looping and localized coil-globule transition. In most prior applications, a single protein or a tightly preassembled protein complex were investigated^33–47^. Sharma demonstrated that digestion of a single compacting agent by Proteinase K reverses compaction^59^. However, many, if not the majority, of DNA-compacting proteins do not function in isolation but rather are part of multi-protein complexes in their *in vivo* context. Once we consider multi-protein complexes, an interesting question arises as to whether the assembly sequence of the multi-protein complex impacts the final DNA configuration. For instance, allostery may demand that the multiprotein complex form prior to binding DNA, or the complex could assemble through step-wise assembly where one member of the complex first binds to DNA and then recruits all others components.

In order to demonstrate that the binding sequence can be of essential importance in multi-protein assemblies, we used two proteins from the telomere protein complex^61,73^. The telomere is a multi-protein assembly that protects the ends of linear chromosomes in eukaryotic cells from recognition as doubled stranded breaks by forming a T-loop. Specifically, we chose the telomeric TRF1 and TIN2 protein, where TRF1 can bind DNA in dimers^74**?**, 75^, while TIN2 cannot bind DNA, but specifically interacts with TRF1^76,77^. Using atomic force microscopy, we previously showed that TRF1 has the ability to compact DNA^65^, and that TIN2 increases the ability of TRF1 to drive the compaction^66^.

In Fig. 5A we demonstrate the nanofluidic implementation of the commonly-performed experiment in which a buffer with premixed TRF1 and TIN2 is added to DNA. DNA is initially observed in a protein-free buffer after it is confined in a nanogroove. DNA is then exposed to the protein solution from the “protein 1” channel in the dark for 5 min, and then observed again to reveal an apparent approximately homogeneous compaction (Fig. 5B). A further 5 min exposure to the protein mixture does not increase the compaction.

**FIG. 5.**
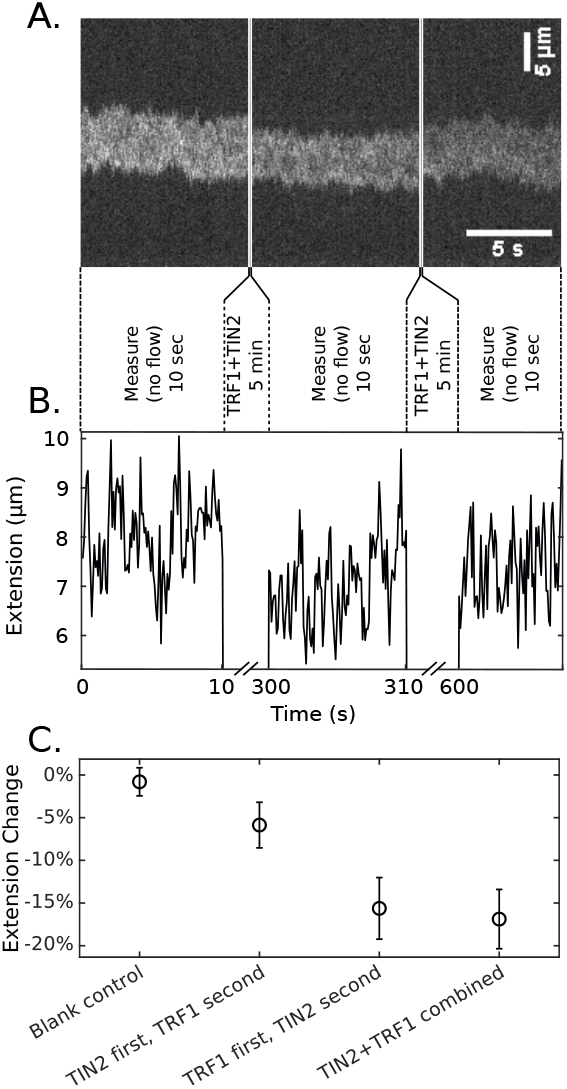
Compaction of ligated telomeric DNA substrates under exposure to 900 nM of TRF1 and TIN2. 5-min exposure steps under constant flow in the nanoslit were followed by 10-sec observation steps without flow. **A**. Kymograph of simultaneous exposure. **B**. Extension of molecule in Panel A. **C**. Analysis determining impact of exposure sequence. Points are median values, and error bars are 2-*σ* values derived from the median absolute deviation.

We executed this experimental procedure with premixed proteins (as shown), by having TRF1 in the first exposure (“protein 1” channel) and TIN2 in the second exposure (“protein 2” channel), by having TIN2 in the first and TRF1 in the second exposure, or by having no protein at all (blank buffer). We quantified the compaction using reported methods^31^, and find strong impact of the incubation sequence (Fig. 5C). In particular, the median telomeric DNA compaction after incubation with TIN2 followed by TRF1, with TRF1 followed by TIN2, and with the TRF1-TIN2 mixture were -6±1%, - 16±2%, and -17±2%, respectively. A larger study comprising TRF1, TIN2, as well as SA1 with a careful exploration of the interaction network has been submitted for review^62^. We believe that the basic idea of exploring the impact of protein exposure sequences will lead to the elucidation of the assembly mechanism of system such as chromatin remodeling complexes that contain a multitude of proteins and interaction points.

### D. Active Enzymatic DNA Processing

Enzymatic processing of DNA is at the heart of its biological function. If this processing is linked to large-scale conformation changes, such double-stranded scission or processive extrusion of a loop, then it can be observed in nanofluidic experiments.

In Fig. 6 we demonstrate the real-time digestion of *λ* - DNA with SmaI endonuclease. We had earlier shown that the requirement of endonucleases to bind divalent metal ion to cleave DNA can be used to localize their reaction in nanochannels^48^. Specifically, digestion of *λ* -DNA with SmaI requires Mg^2+^ and results in four easily observable fragments of length 19.4 kbp, 12.2 kbp, 8.3 kbp, and 8.6 kbp. DNA was pushed into nanogrooves in the Mg^2+^-free buffer, and an otherwise identical buffer that contained 4 mM of Mg^2+^ was flushed in from the “protein 1” channel as soon a DNA had settled into nanogrooves. As the buffer containing Mg^2+^ reaches the confined DNA, the cutting of DNA starts. Four distinct fragments consistent with the sequence of *λ* -DNA are observed Fig. 6.

**FIG. 6.**
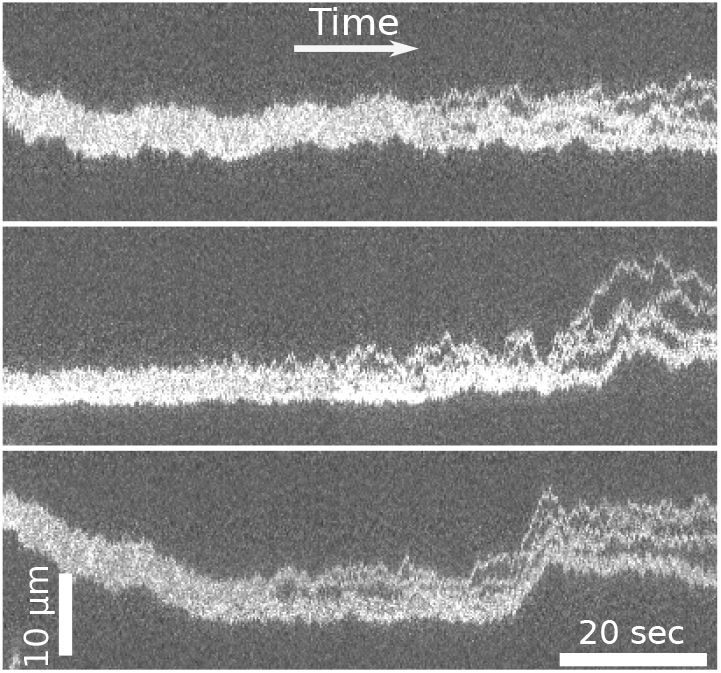
Kymographs showing restriction mapping of *λ* -DNA with the SmaI endonuclease. DNA was brought into the channel in absence of Mg^2+^ and the Mg^2+^-carrying buffer was flushed over it.

While the endonuclease uses a protein which utilizes the energy inherent to the substrate to drive its catalytic action, the majority of DNA-modifying enzymes use turnover of a co-factor such as ATP hydrolysis to bias the chemical equilibrium towards the final state. We have previously shown the activity of T4 DNA ligase in nanochannels^78^, but could not report a conformation change of the DNA at that point. Here we demonstrate the processive ATPase-dependent DNA loop extrusion by cohesin-NIPBL, which has previously been established using tethered-end DNA assays^79,80^. However, in these publications DNA end were anchored and the loop junction and DNA loop extrusion was observed under flow. Loop extrusion could only proceed under tension, or without direct observation.

In Fig. 7 we demonstrate the real-time loop extension using NIPBL-cohesin. The protein was loaded onto T4 phage DNA, and DNA was brought into a Y-junction identical to the ones used for flow manipulation. If the molecule contained a loop, then the DNA loop anchor point typically at first was not located at the 3-channel junction of the Y, but rather in the stem or one of the branches as evidenced by steps of the brightness along the channel axis. We brought the DNA loop anchor to the junction point, and ceased all flow. Using a wild-type NIPBL-cohesin, we were then able to observe DNA loop mobility and specifically loop extrusion (Fig. 7A.-C.), while static loops also were observable. In a ATPase-deficient mutant, only static loops were present (Fig. 7D. and E). The designation of dynamic and extruding loops versus static loops does not imply that the visible length in the stem and branches was invariant and not fluctuating for a static loop. Rather, it implies that linear fits to the slopes of the lengths were small (≤ 0.15 µm/s), while the active loop extrusion resulted in slopes ≥ 0.4 µm/s.

**FIG. 7.**
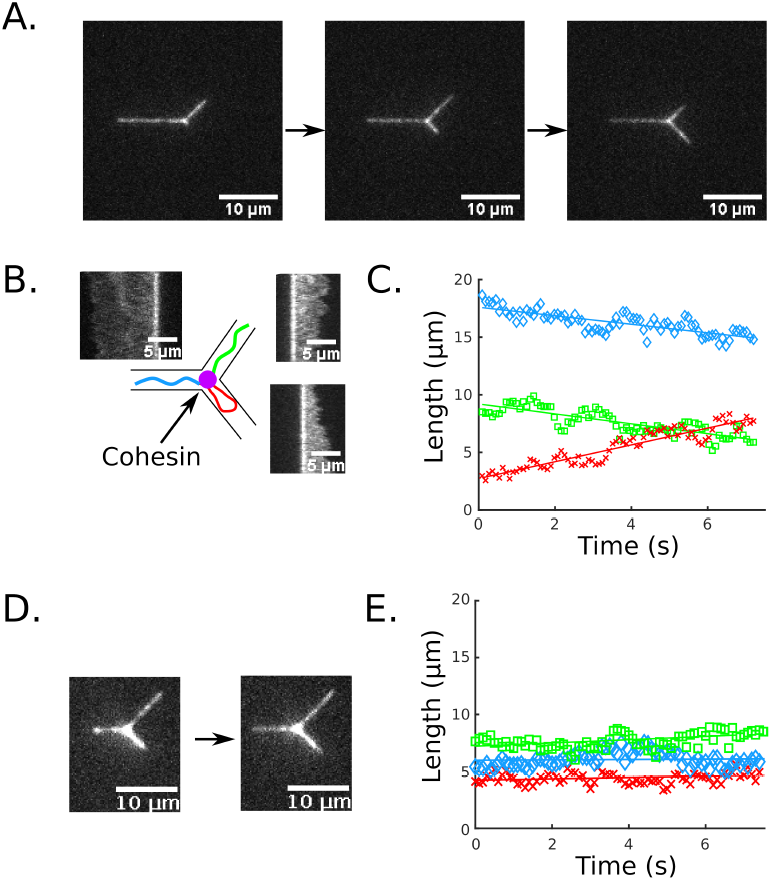
Loop extrusion by cohesin-NIPBL. **A**. Still frames from molecule undergoing loop extrusion under a no-flow condition. The hernia is to the bottom right branch. **B**. Kymographs covering 7.2 sec of loop extrusion in panel A., limited by subsequent breakup of molecule under oxidative stress. Kymographs are located adjacent to the channel that they occupy, and the loop is located in the bottom right branch. Cohesin is indicated as a purple in the schematic, and the DNA molecule is locally colored according to the channel in which the respective stretch resides. **C**. Lengths of DNA in the channels adjacent to the 3-way junction (blue ♢ single-occupied stem, green □ singly occupied top branch, red doubly occupied looped in bottom branch). **D**. ATPase-deficient mutant shows no extrusion activity. The loop, which is possibly nested is located in the bottom right branch. **E**. Lengths of legs from data underlying panel D. using the same labeling scheme as in panel C..)

## IV. DISCUSSION AND CONCLUSION

In conclusion, we have demonstrated how a device following the mixed 2d-1d confinement paradigm can perform complex DNA configuration manipulations as well as programmable exposure patterns to proteins and their co-factors under continuous fluorescence imaging. Specifically, we expand the range of capabilities to testing the impact that the exposure sequence of a DNA-compacting two-protein system has, we demonstrate the controlled manipulation of DNA hernias and multi-DNA end-contact at Y-junctions, and we observe the active loop extrusion by a protein at such a Y-junction. The expansion of the capabilities of the mixed 2d-1d-paradigm points to a broad set of possible studies that can be performed using the same device. For instance, we explore the interactions of a three-protein system that has been proposed as responsible for telomere cohesion during replication^81^ in an simultaneously submitted manuscript^62^. A large class of sequentially-assembled protein constructs such as the complexes regulating transcription naturally lend themselves to the same method^82^. Similarly, the interplay of condensin and cohesin with DNA poses a large number of open questions^83^ for which interrogation of DNA loop extrusion in nanochannels can play an important role.

## ACKNOWLEDGMENTS

This work was supported by the National Institutes of Health (R01GM107559 to H. W., R. R., R01GM123246 to H. W., R. R., P. L. O., P30 ES025128 Pilot Project Grant to H. W. and P. K. through the Center for Human Health and the Environment at NCSU, and R01CA207342 to P.L.O.). This work was supported by the National Natural Science Foundation of China (Project 32130053 to H.Y.).

This work was performed in part at the Analytical Instrumentation Facility (AIF) at North Carolina State University and the NCSU Nanofabrication Facility (NNF), which are supported by the State of North Carolina and the National Science Foundation (award number ECCS-2025064). AIF and NNF are members of the North Carolina Research Triangle Nanotechnology Network (RTNN), a site in the National Nanotechnology Coordinated Infrastructure (NNCI).

## DATA AVAILABILITY STATEMENT

The data that support the findings of this study are available from the corresponding author upon reasonable request.

## Notes

### Competing Interest Statement

The authors have declared no competing interest.

